# Shared Genetic Architecture between Muscle and Bone: Identification and Functional Implications of *EPDR1*, *PKDCC*, and *SPTBN1*

**DOI:** 10.1101/2023.05.14.540743

**Authors:** Jongyun Jung, Qing Wu

**Author notes:** Corresponding author (QW).

## Abstract

Recent studies suggest a shared genetic architecture between muscle and bone, yet the underlying molecular mechanisms remain elusive. This study aims to identify the functionally annotated genes with shared genetic architecture between muscle and bone using the most up-to-date genome-wide association study (GWAS) summary statistics from bone mineral density (BMD) and fracture-related genetic variants. We employed an advanced statistical functional mapping method to investigate shared genetic architecture between muscle and bone, focusing on genes highly expressed in muscle tissue. Our analysis identified three genes, *EPDR1, PKDCC*, and *SPTBN1*, highly expressed in muscle tissue and previously unlinked to bone metabolism. About 90% and 85% of filtered Single-Nucleotide Polymorphisms were located in the intronic and intergenic regions for the threshold at *P* ≤ 5 × 10^−8^ and *P* ≤ 5 × 10^−100^, respectively. *EPDR1* was highly expressed in multiple tissues, including muscle, adrenal gland, blood vessels, and thyroid. *SPTBN1* was highly expressed in all 30 tissue types except blood, while *PKDCC* was highly expressed in all 30 tissue types except the brain, pancreas, and skin. Our study provides a framework for using GWAS findings to highlight functional evidence of crosstalk between multiple tissues based on shared genetic architecture between muscle and bone. Further research should focus on functional validation, multi-omics data integration, gene-environment interactions, and clinical relevance in musculoskeletal disorders.

**Author Summary:** Osteoporotic fractures in the aging population pose a significant health concern. They are often attributed to decreased bone strength and muscle loss. However, the underlying molecular connections between bone and muscle are not well understood. This lack of knowledge persists despite recent genetic discoveries linking certain genetic variants to bone mineral density and fracture risk. Our study aimed to uncover genes that share genetic architecture between muscle and bone. We utilized state-of-the-art statistical methods and the most recent genetic data related to bone mineral density and fractures. Our focus was on genes that are highly active in muscle tissue. Our investigation identified three new genes - *EPDR1, PKDCC*, and *SPTBN1* - which are highly active in muscle tissue and influence bone health. These discoveries offer fresh insights into the interconnected genetic makeup of bone and muscle. Our work not only uncovers potential targets for therapeutic strategies to enhance bone and muscle strength but also provides a blueprint for identifying shared genetic structures across multiple tissues. This research represents a crucial step forward in our understanding of the interplay between our muscles and bones at a genetic level.

## Introduction

Osteoporosis, a disease characterized by low bone mineral density (BMD), decreased bone strength, and increased fracture risk, poses a significant challenge for the aging population [1,2]. Concurrent muscle loss exacerbates this issue, further contributing to weakened bone strength and heightened susceptibility to osteoporotic fractures [3,4]. Together, osteoporosis and muscle loss represent significant risk factors for fractures in older adults [5,6].

Adequately developed muscles play a crucial role in protecting osteoporosis patients from fractures by reducing the likelihood of falls and shielding the body from osteoporotic fractures [7]. However, aging is commonly associated with the loss of skeletal muscle, which, in turn, can lead to declines in physical functioning among older adults [5,6]. Despite numerous genetic studies investigating BMD or fracture-related outcomes, the genetic link between bone and muscle remains underexplored. The exact biological pathway through which bone influences muscle is still unknown [3,8,9].

Over the past decade, genome-wide association studies (GWAS) have identified multiple genetic variants linked to BMD or fractures [10,11]. A recent study of 426,824 UK Biobank participants discovered 518 loci associated with ultrasound-derived heel BMD [10]. However, GWAS has had limited success in pinpointing causal genes, with the majority of findings located in non-coding or intergenic regions [12–14]. Functional mapping is commonly used in translational research to select and prioritize genetic variants likely to be functional and quantify the strength of evidence in the GWAS [15]. In this study, we employ FUMA, a comprehensive functional mapping method that utilizes 18 biological data repositories and tools to annotate and prioritize GWAS findings [16].

Our study aims to investigate the functional consequences of BMD or fracture-related GWAS findings at the muscle tissue level, providing novel insights into the genetic architecture connecting muscle and bone. We hypothesize that identifying BMD or fracture-related GWAS findings associated with muscle loss will shed light on the genetic interplay between these two tissues and contribute to a deeper understanding of their relationship in the context of osteoporosis. Genetic variants underlying bone and muscle, identified using GWAS and statistical functional mapping methods, will provide a novel insight into targets to stratify patients for risk and develop new therapeutic strategies.

## Methods

### Genome-Wide Association Study (GWAS) Summary dataset and Pre-processing

We utilized the comprehensive GWAS dataset from Morris et al. [10], which investigated genetic variants of bone mineral density (BMD) estimated by heel quantitative ultrasound (eBMD) in 426,824 UK Biobank participants. The study identified 518 genome-wide BMD-significant loci and 13 bone fracture loci. We applied two P-value thresholds (*P* ≤ 5 × 10^−8^ and *P* ≤ 5 × 10^−100^) to filter significant SNPs from the GWAS summary dataset. Using different P-value thresholds, we examined whether bone and muscle tissues shared different genetic architectures. A P-value threshold of *P* ≤ 5 × 10^−100^ is widely used to identify associations between common genetic variants and traits. A more stringent P-value threshold of *P* ≤ 5 × 10^−100^ was utilized to investigate whether rare variants contribute to the shared genetic architecture between bone and muscle. The GWAS summary statistics (UK Biobank eBMD and Fracture GWAS Data Release 2018) were obtained from the GEnetic Factor for Osteoporosis (GEFOS) consortium website [17]. Data pre-processing was performed using R version 3.6.1 software (The R Foundation)) [18].

### Functional mapping with FUMA

We employed FUMA [16], a functional mapping and annotation tool, to prioritize the most likely causal single-nucleotide polymorphisms (SNPs) and genes using information from 18 biological data repositories and tools. FUMA’s core functions, SNP2GENE and GENE2FUNC processes, were used to annotate SNPs with biological functionality, map them to genes based on positional, eQTL, and chromatin interaction information, and gain insight into the putative biological mechanisms of prioritized genes.

### Characterization of Genomic Risk Loci and Gene Annotation

Independent and significant SNPs were identified using the 1000 Genomes Project reference panel (*r*^2^ < 0.6). Independent lead SNPs were defined based on an *r*^2^ < 0.1. Genomic loci of interest were grouped if LD blocks of independent and significant SNPs were closely located (<250 kb). Gene annotation was performed using ANNOVAR [19] and Ensemble genes (build 85).

### Gene-based and Gene-set Enrichment Analysis

Gene-based GWAS analysis was conducted with MAGMA 1.6 [20] using default settings implemented in FUMA [16]. The gene-based P-values were employed for gene-set enrichment analyses, where gene expression values for 53 specific tissue types were obtained from GTEx [16]. A significance level of *P* ≤ 5 × 10^−10^ and *P* ≤ 5 × 10^−100^ corresponded to thresholds of 1.69 × 10^−5^ and 1.67 × 10^−3^, respectively. Functional annotation of GWAS results was performed using FUMA [16]. Normalized gene expression data (reads per kilobase per million, RPKM) for 53 tissue types were obtained from GTEx portal v6, yielding transcripts for 28,520 genes, with 22,146 mapped to the Entrez gene ID.

## Results

### Summary of Genome-Wide Association Study Statistics

We analyzed previously published summary statistics from a genome-wide association study (GWAS) on bone mineral density estimated from quantitative heel ultrasounds (eBMD) [10], which included genotyping and imputed data for up to 426,824 participants in the U.K. Biobank study available from the GEnetic Factor for OSteoporosis consortium (GEFOS) website [17] (**Fig 1A**). We performed two primary analyses using FUMA: characterization of significant hits and genome-wide analysis (**Fig 1B**).

**Fig 1.**
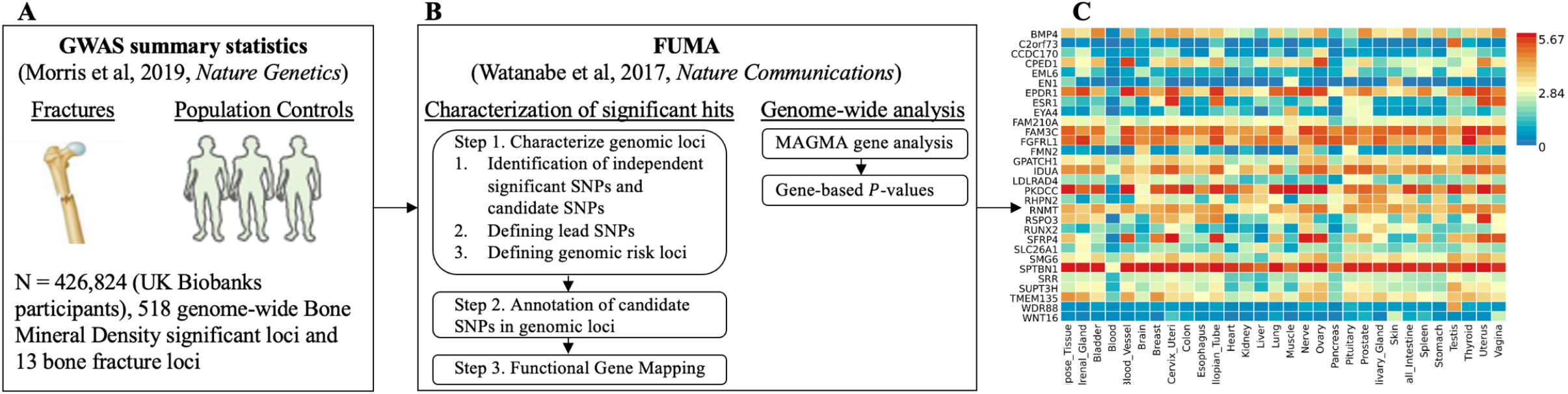
Study flow. **A**) Genome-Wide Association Study (GWAS) summary statistics of genetic association with Bone Mineral Density (BMD) or fractures for 13,753,401 Single Nucleotide Polymorphisms (SNPs) from a study estimated in 426,824 UK Biobank participants study [10]. **B**) FUMA [16] is a web-based platform utilizing information from multiple biological resources for functional annotation of GWAS results and prioritizing the most likely causal SNPs using information from 18 biological data repositories, including the Genotype-Tissue Expression (GTEx) data. Characterization of significant hits and Genome-wide analysis were conducted with FUMA. **C**) Heatmap shows the highly expressed genes in multiple tissues of GTEx data.

The eBMD GWAS summary statistics contained data for 13,753,401 SNPs. We filtered the summary statistics using two thresholds: *P* ≤ 5 × 10^−8^ and *P* ≤ 5 × 10^−100^. Thesethresholds yielded 103,155 and 1,724 candidate GWAS-tagged SNPs, mapping to 2,955 and 30 genes, respectively (**Table 1**). Approximately 90% and 85% of filtered SNPs were located in intronic and intergenic regions at *P* ≤ 5 × 10^−8^ and *P* ≤ 5 × 10^−100^ thresholds, respectively (**Fig 2**). The summary per genomic risk locus plot revealed that genomic loci 6:44683838-45404170 had the largest size (>700 kb), followed by 2:119014660-119634677 (>600 kb) and 2:54616729-55003484 (>400 kb) (**S1 Fig**). Over 550 SNPs were found in the genomic loci — 7:120703929-121098222.

**Table 1.** The summary of Genome-Wide Association Study (GWAS) Summary Statistics for Single-Nucleotide Polymorphisms (SNPs) and mapped genes. Two different *P*-value thresholds were used to filter the Genome-Wide Association Study (GWAS) Summary Statistics GWAS summary statistics.

| | $P \leq 5 \times 10^{-8}$ | $P \leq 5 \times 10^{-100}$ |
| --- | --- | --- |
| Number of candidates GWAS tagged SNPs | 103,155 | 1,724 |
| Number of lead SNPs | 2,410 | 40 |
| Number of genomic risk loci | 600 | 25 |
| Number of mapped genes | 2,955 | 30 |

**Fig 2.**
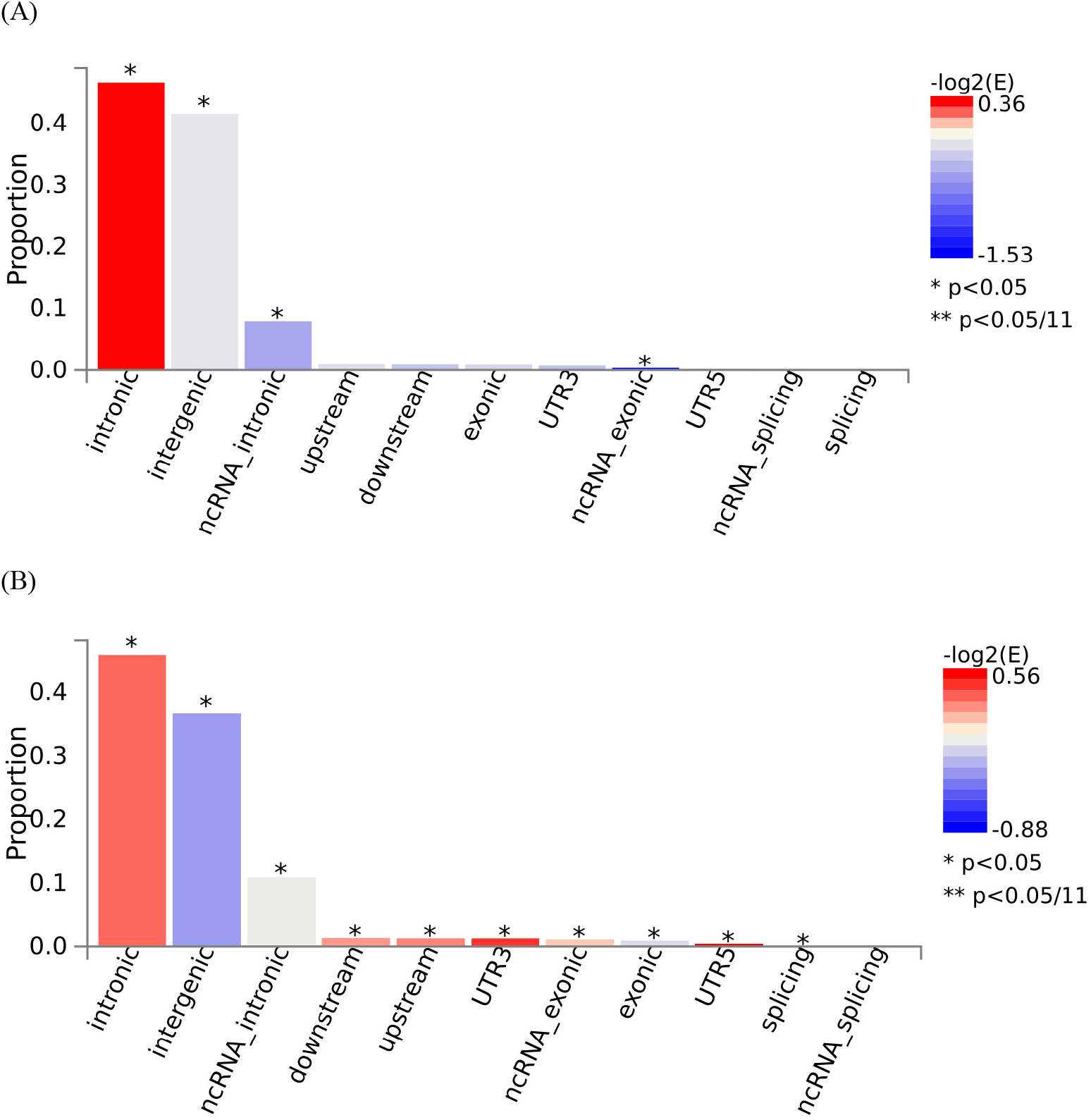
Functional consequences of Single Nucleotide Polymorphisms (SNPs) with two different thresholds. Two different P-value thresholds **(A)** *P* ≤ 5 × 10^−8^and (B) *P* ≤ 5 × 10^−100^ were used to filter the Genome-Wide Association Study (GWAS) Summary Statistics.

**S1 Fig.**
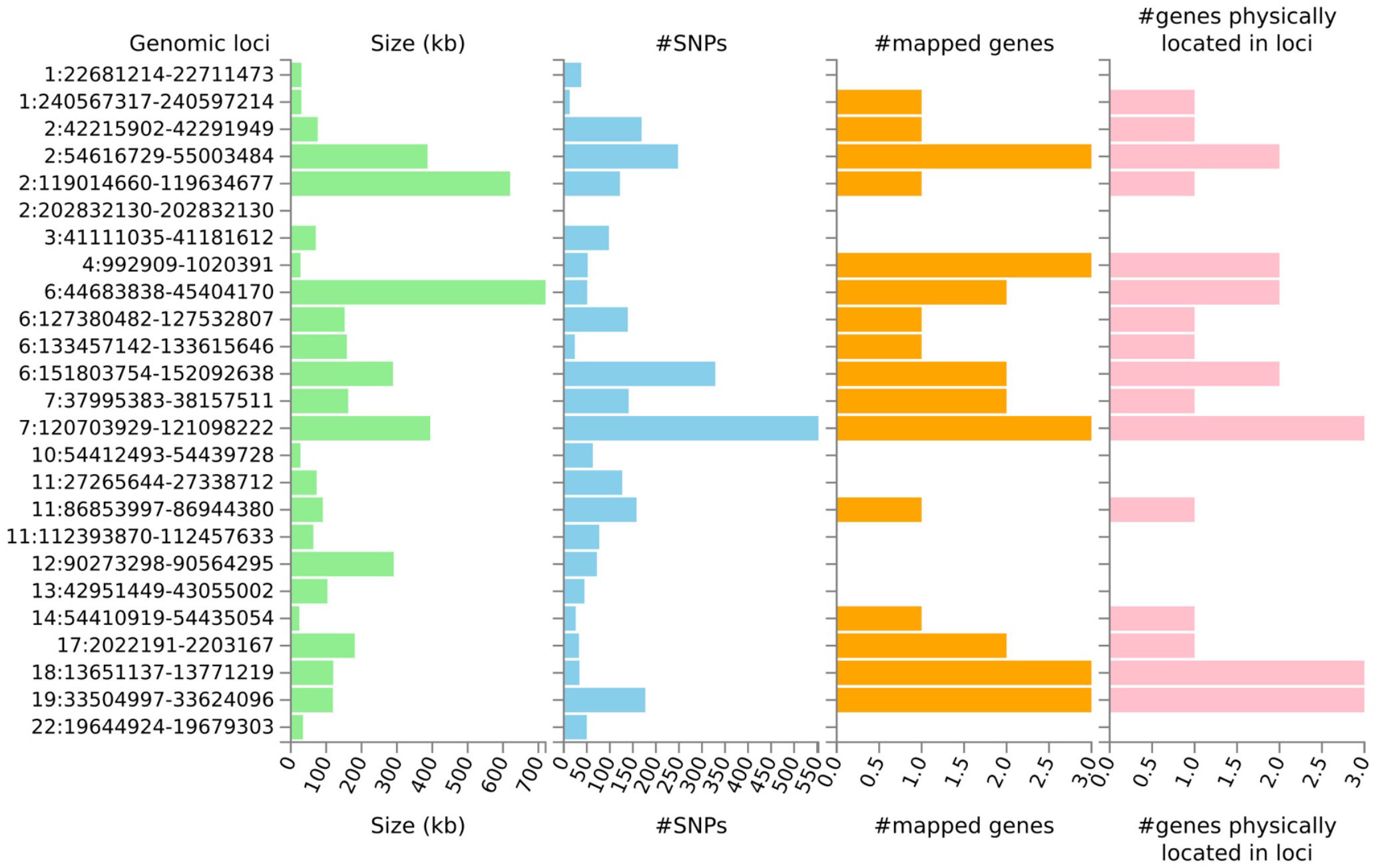
The summary per genomic risk locus. Note that genomic loci could contain more than one independent lead Single-Nucleotide Polymorphisms (SNPs).

Manhattan plots at the SNP level for both thresholds indicated that chromosomes 2, 6, 7, 10, and 22 contained the most significant SNPs (**Fig 3**). At the gene level, multiple genes were identified, including *NKX1-1, ZNF800, EML6, EN1*, and *LDLRAD4* (**Fig 4**).

**Fig 3.**
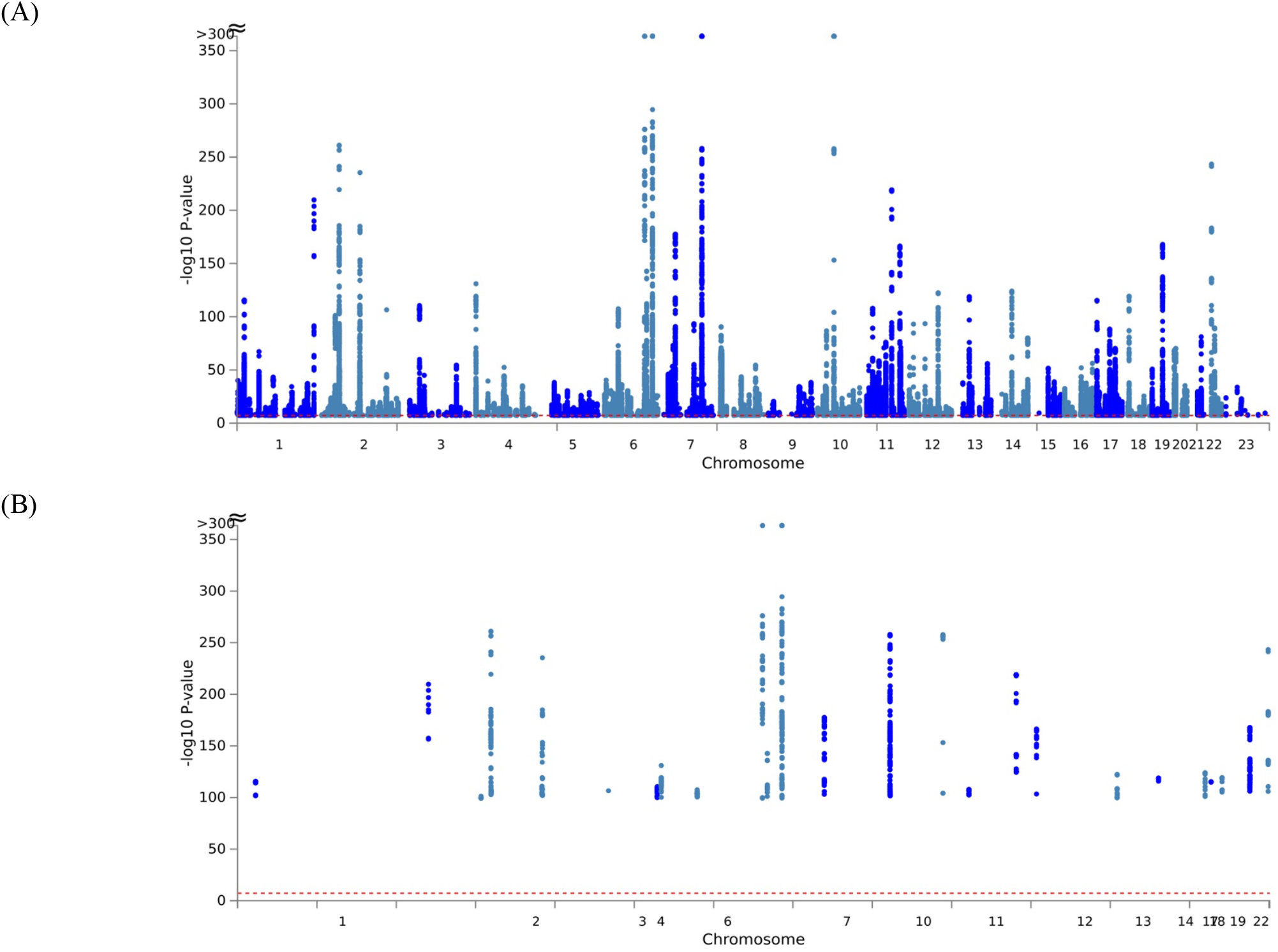
Manhattan Plot at Single Nucleotide Polymorphisms (SNPs) level. (A) *P* ≤ 5 × 10^−8^ and (B) *P* ≤ 5 × 10^−100^ were used to filter the Genome-Wide Association Study (GWAS) Summary Statistics.

**Fig 4.**
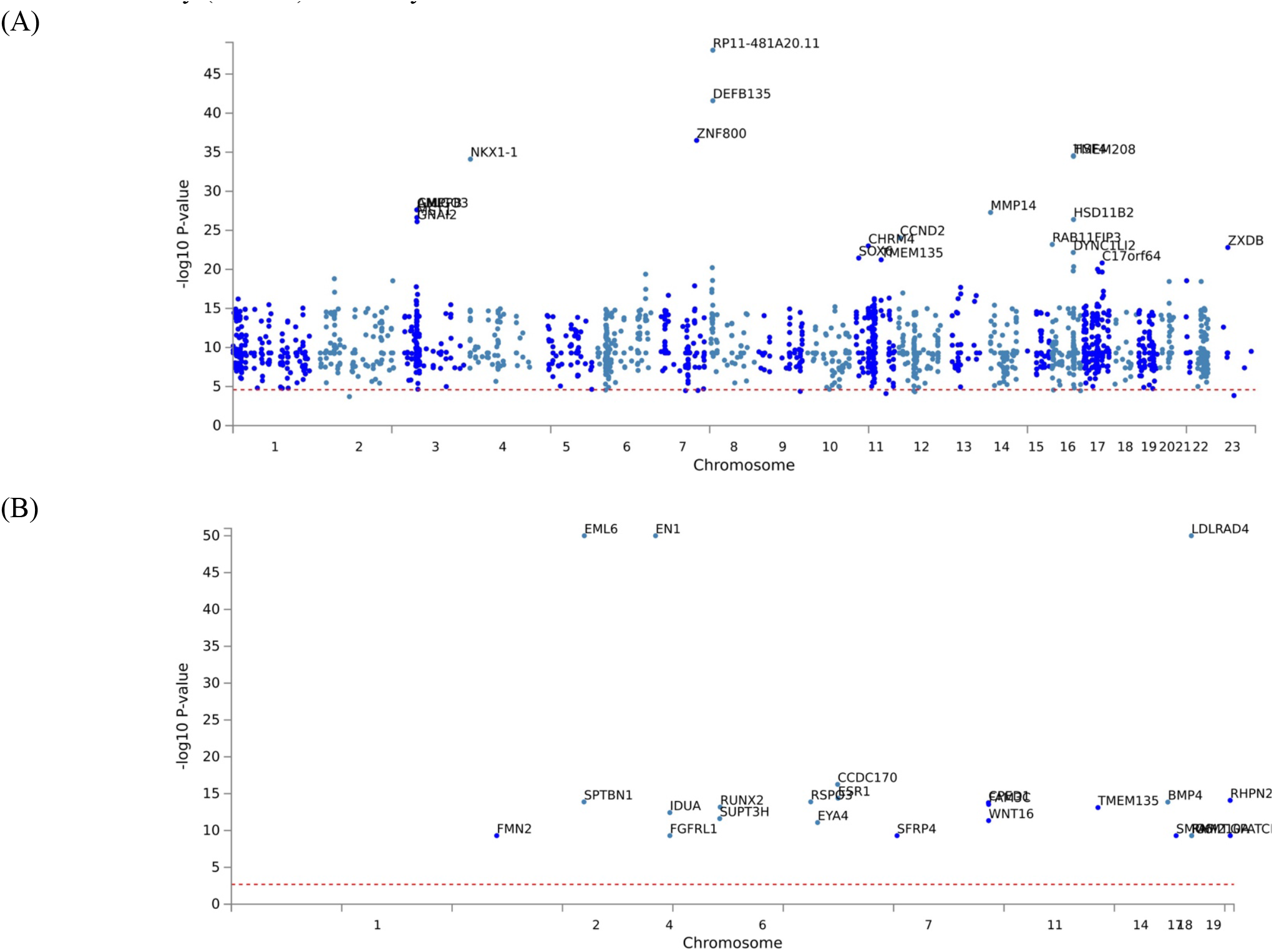
Manhattan Plot at the gene level. (A) *P* ≤ 5 × 10^−8^ and (B) *P* ≤ 5 × 10^−100^ were used to filter the Genome-Wide Association Study (GWAS) Summary Statistics.

### Gene Expression Analysis at the Tissue Level

*EPDR1, PKDCC*, and *SPTBN1* genes showed high expression levels in muscle tissue (**Fig 5**). *EPDR1* was also highly expressed in other tissues, such as the adrenal gland, blood vessel, cervix uteri, fallopian tube, nerve, ovary, thyroid, and uterus. *SPTBN1* demonstrated high expression across all 30 tissue types except blood, while *PKDCC* was highly expressed in all tissue types except the brain, pancreas, and skin. Tissue enrichment analysis using MAGMA results indicated that the most enriched tissue was the mammary breast tissue at the *P* ≤ 5 × 10^−100^ threshold (**Fig 6**), and several other tissues, including fallopian tube, ovary, kidney cortex, kidney medulla, breast mammary, cervix endocervix, and adipose visceral omentum, were enriched at the *P* ≤ 5 × 10^−100^ threshold (**S2 Fig**). Differentially expressed genes (DEG) in 30 major tissues from GTEx data showed up-regulated DEG in breast and prostate tissues at the 5 × 10^−100^ threshold (**Fig 6**). However, no significant DES was observed at the *P* ≤ 5 × 10^−8^ threshold (**S3 Fig**).

**Fig 5.**
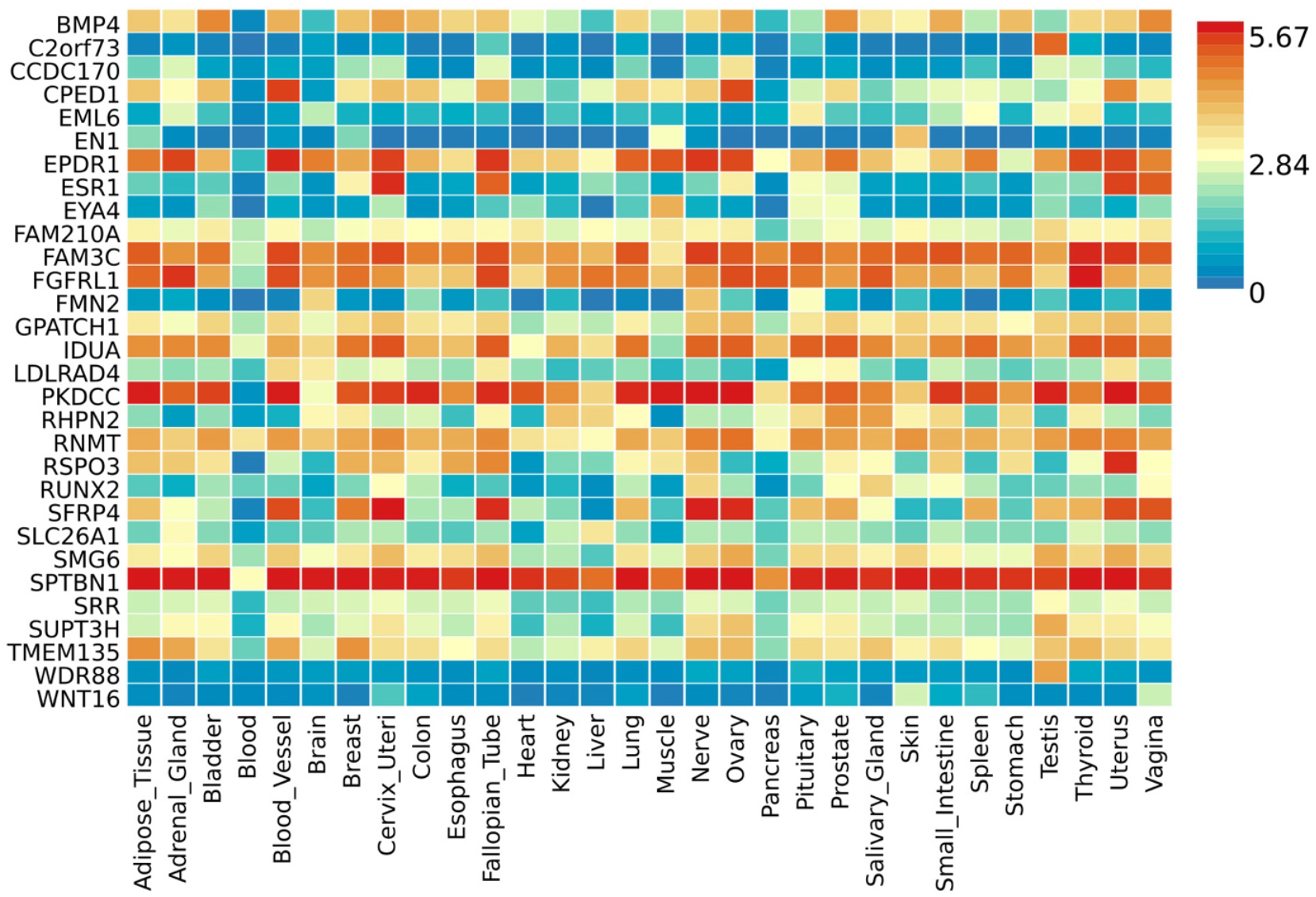
Gene expression heatmap with the GTEx v8 30 tissue types. A threshold *P* ≤ 5 × 10^−100^was used to map the genes.

**Fig 6.**
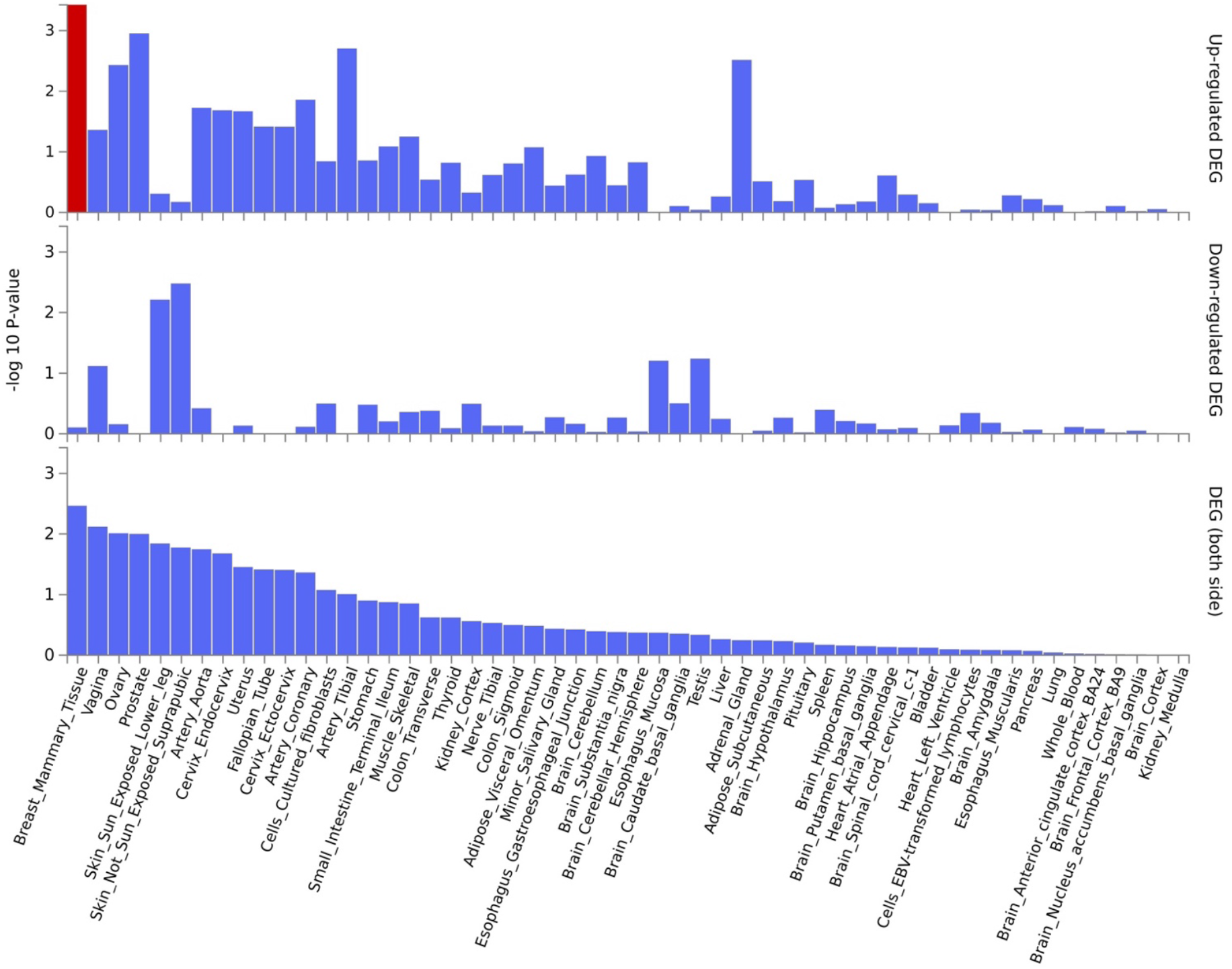
Tissue enrichment analysis using MAGMA [20]. The most enriched tissue is the mammary breast tissue. Significantly enriched DEG sets (*P*_bon;_ < 0.05) are highlighted in red. A threshold *P* ≤ 5 × 10^−100^ was used to map the genes.

**Figure 7.**
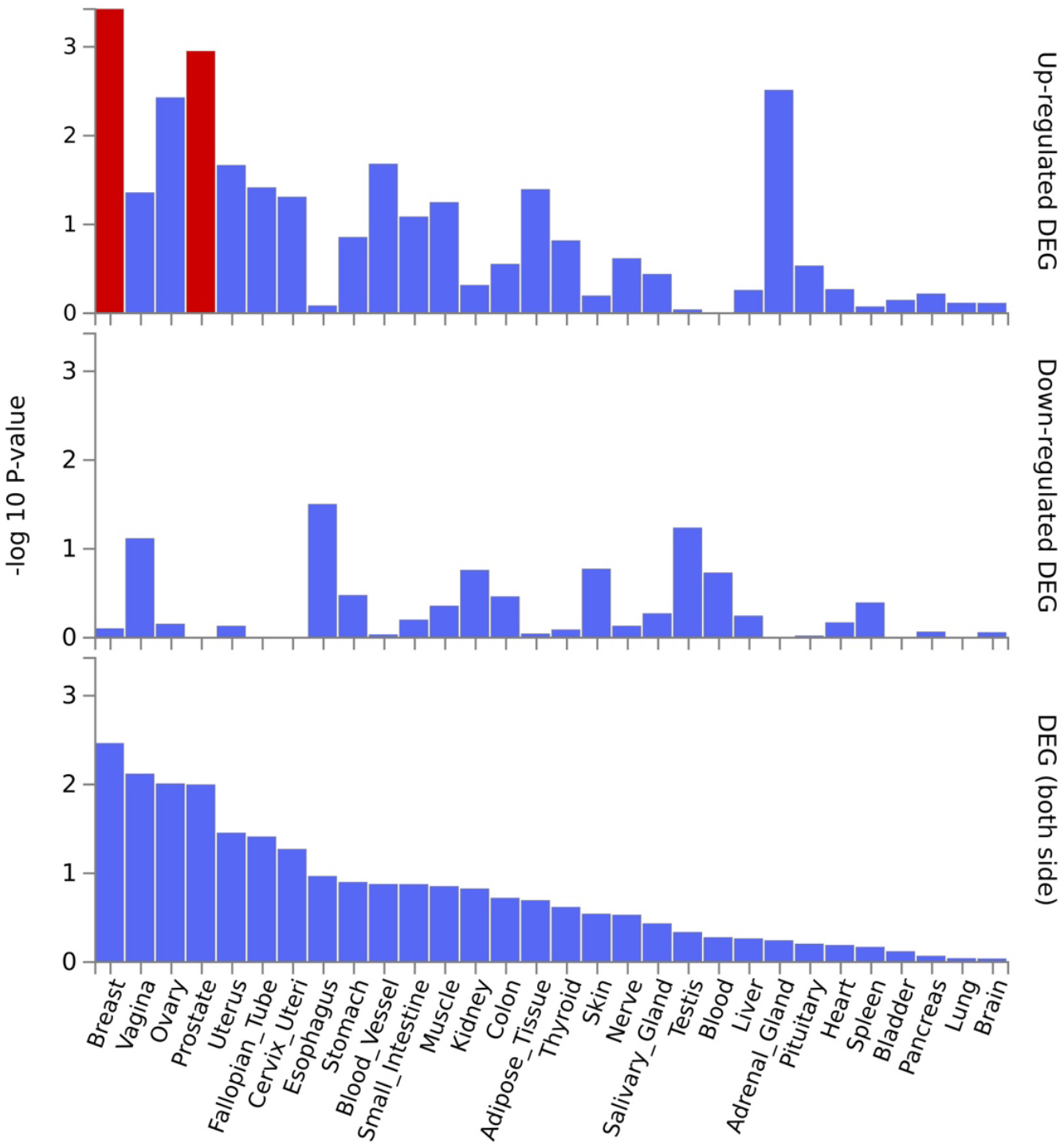
Differentially expressed genes in 30 major tissues in the GTEx data. A threshold *P* ≤ 5 × 10^−100^ was used to map the genes.

**S2 Fig.**
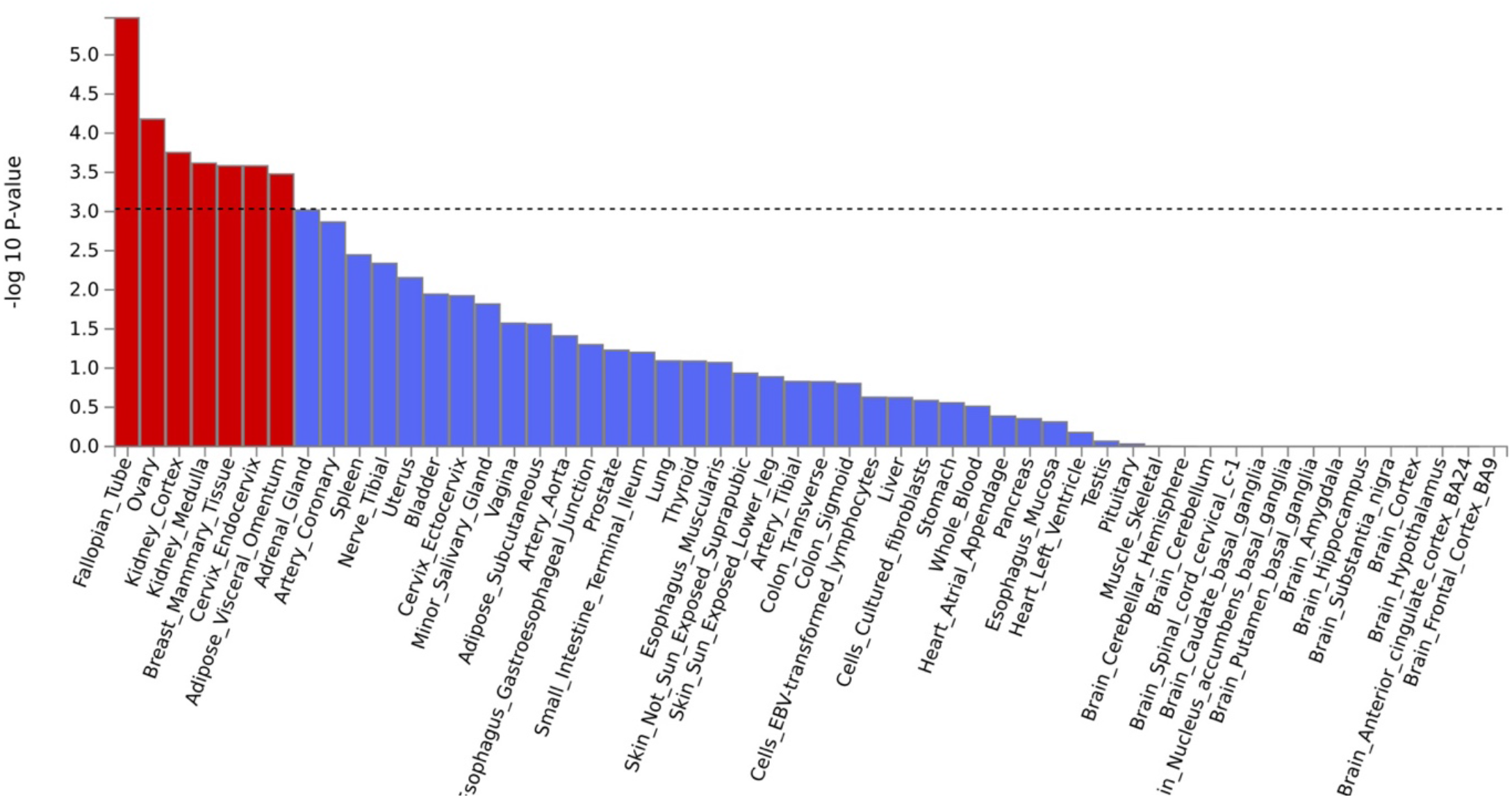
Tissue enrichment analysis using MAGMA [20]. The most enriched tissue is the fallopian tube. Significantly enriched DEG sets (*P*_bon_ < 0.05) are highlighted in red. A threshold *P* ≤ 5 × 10^!”^ was used to map the genes.

**S3 Fig.**
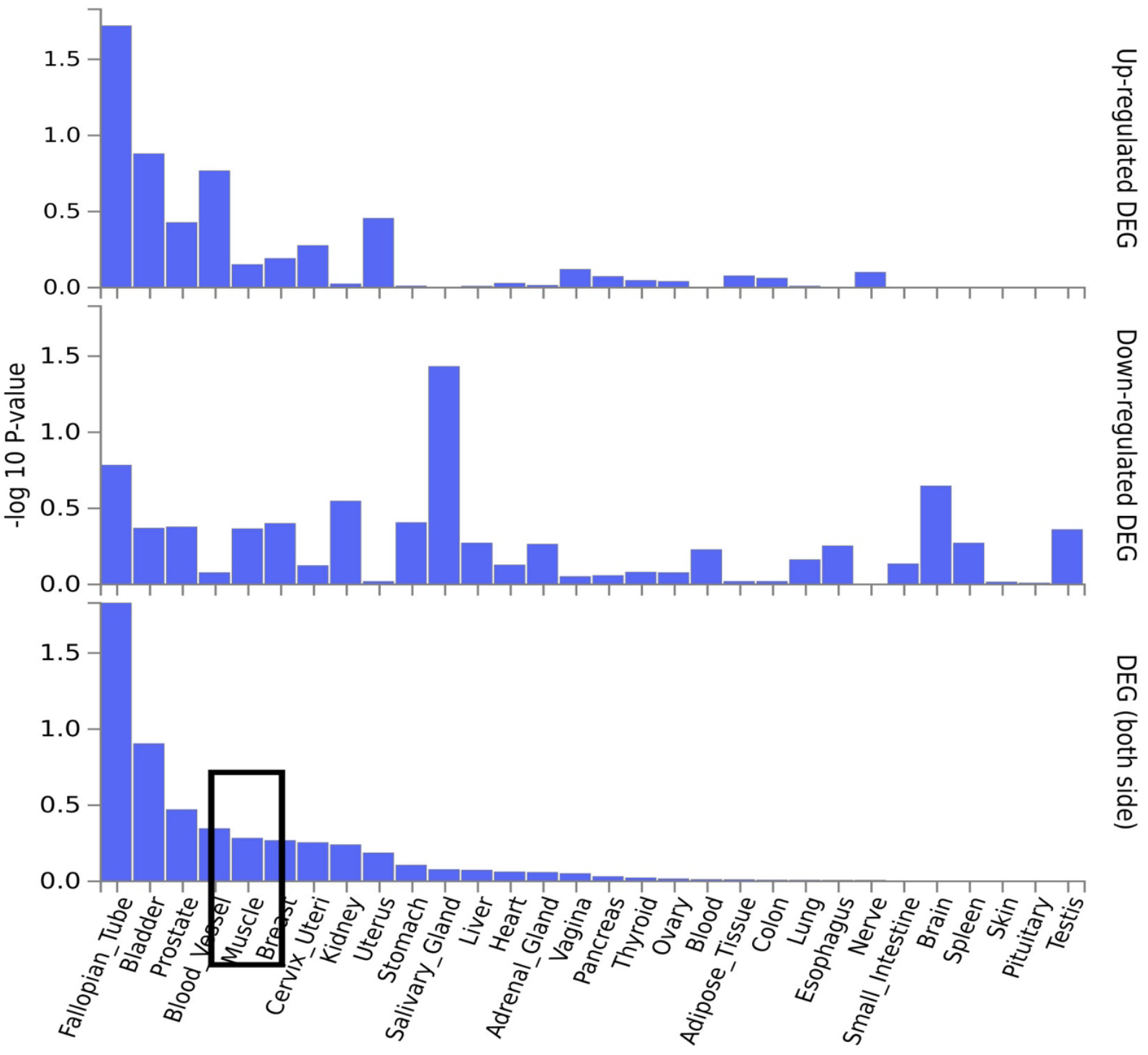
Differentially expressed genes in 30 major tissues in the GTEx data. A threshold *P* ≤ 5 × 10^!”^ was used to map the genes.

## Discussion

Our study identified three novel genes — *EPDR1, PKDCC*, and *SPTBN1* — highly expressed in muscle tissue using the most updated GWAS summary statistics from BMD and fracture-related genetic variants, coupled with advanced statistical fine-mapping methods. These genes were not previously associated with both muscle and bone tissues, providing new insights into the shared genetic architecture of these systems. Our findings support the existence of pleiotropy between muscle and bone, as previously suggested [21].

*EPDR1* has been shown to be highly expressed in the brain and detected in other tissues, such as muscle, heart, and extracellular fluids [22]. *SPTBN1* was linked to BMD in a previous study [23], and our results suggest that it plays a crucial role in bone metabolism and maintaining the shared genetic architecture between bone and muscle. *PKDCC* has not yet been implicated in BMD or fracture-related outcomes. Our study contributes to understanding biological pathways affecting muscle and bone, as previously described [9,21,24], and complements earlier GWAS findings [23,25].

A noteworthy observation in our analysis is that 90% and 85% of filtered SNPs were located in intronic and intergenic regions for the thresholds at *P* ≤ 5 × 10^−8^ and *P* ≤ 5 × 10^−100^, respectively. Although GWAS have identified numerous genetic variants associated with BMD and fracture, their applicability to functional studies remains limited. Consequently, additional genetic variants might be relevant to the relationship between muscle and bone genetic architecture that were not captured in our analysis.

Our findings should be interpreted in the context of certain limitations. First, the GWAS summary statistics used in our study were derived from a White population; future research should validate our findings in other racial/ethnic people. Second, our analysis assumes that GWAS findings from fractures or BMD are related to muscle tissue, but this approach may not capture all relationships between muscle and bone genetic architecture. Trajanoska et al. [9] highlighted the influence of mitochondrial genetics on bones and muscles, which standard GWAS cannot reliably assess due to the sparse number of mitochondrial markers on genotyping arrays and difficulties in quantifying mitochondrial heteroplasmy. Moreover, many haplotype blocks containing GWAS SNPs do not overlap with regions of known function and remain classified as intronic or intergenic [26].

In conclusion, our study has identified three new genes — *EPDR1, PKDCC*, and *SPTBN1* — highly expressed in muscle tissue, providing novel insights into the shared genetic architecture between muscle and bone. Furthermore, we offer a framework for utilizing GWAS findings to emphasize functional evidence of crosstalk between multiple tissues based on other genetic architecture. Future research should explore the implications of these findings and investigate potential applications in developing therapeutic strategies targeting both muscle and bone health.

Further research directions to build on our results include: 1) integrating multi-omics data, such as transcriptomics, proteomics, and epigenomics, to uncover the molecular mechanisms linking the identified genes to muscle and bone tissues, 2) validating our novel findings in animal experiments, 3) examining associations between *EPDR1, PKDCC*, and *SPTBN1* expression and the development or progression of musculoskeletal disorders in longitudinal cohort studies to establish their clinical relevance, and 4) exploring novel therapeutic strategies targeting the products or regulatory elements of the identified genes to prevent or treat musculoskeletal disorders. By pursuing these research avenues, we aim to achieve a more comprehensive understanding of the shared genetic architecture between muscle and bone, ultimately benefiting the development of therapeutic interventions targeting both tissues.

## Author Contribution

Conceptualization: Jongyun Jung. Formal analysis: Jongyun Jung. Methodology: Jongyun Jung. Supervision: Qing Wu. Visualization: Jongyun Jung. Writing –original draft: Jongyun Jung.

Writing –review & editing: Jongyun Jung, Qing Wu.

## Data Sharing

The summary statistics used in the current study are available from the GEFOS Consortium at http://www.gefos.org/?q=content/data-release-2018. FUMA, an integrative web-based platform, is freely available at https://fuma.ctglab.nl/.

